# A kinetic model for fluorescence microscopy experiments in disordered media that contains binding sites and obstacles

**DOI:** 10.1101/302422

**Authors:** V.P. Shkilev

## Abstract

A model is proposed that describes the diffusion of molecules in a disordered medium with binding sites (traps) and obstacles (barriers). The equations of the model are obtained using the subordination method. As the parent process, random walks on a disordered lattice are taken, described by the random barriers model. As the leading process, the renewal process that corresponds to the multiple-trapping model is taken. Theoretical expressions are derived for the curves obtained in experiments using fluorescence microscopy (FRAP, FCS and SPT). Generalizations of the model are proposed, allowing to take into account correlations in the mutual arrangement of traps and barriers. The model can be used to find parameters characterizing the diffusion and binding properties of biomolecules in living cells.

## 1 Introduction

Fluorescence recovery after photobleaching (FRAP), fluorescence correlation spectroscopy (FCS) and single particle tracking (SPT) are experimental methods based on the use of fluorescence microscopy. They are widely used in the study of diffusion and binding properties of biomolecules in living cells. The essence of the FRAP experiment is the photobleaching of fluorescing molecules in a small volume and the observation of the rate of fluorescence penetration into this volume. The FCS experiment consists of observing fluctuations in the intensity of fluorescence in a small volume arising from the random wanderings of fluorescing molecules. The SPT experiment directly traces the trajectory of an individual molecule.

For the theoretical description of fluorescence microscopy experiments, the multiple-trapping model is often used (1-4) (In biophysical literature this model has no special name. We use the name adopted in statistical physics.). The equations of this model have the following form:

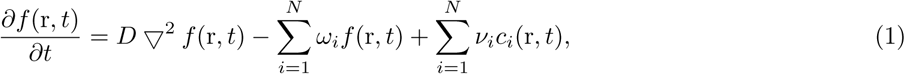

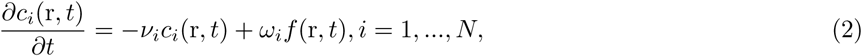

where *D* is the diffusion constant; ∇^2^ is the Laplacian operator; *ω_i_* and *ν_i_* are, respectively, the association and dissociation rates. Variables *f* (r, *t*) and *c_i_*(r, *t*) have different meanings in different experiments. In the FRAP experiment *f* (r, *t*) is the concentration of the unbound freely diffusing molecules and *c_i_*(r, *t*) is the concentration of the bound molecules at binding sites of *i*th type; in the FCS experiment *f* (*r*, *t*) and *c_i_*(r, *t*) are the fluctuations of the concentrations of the unbound and bound molecules, respectively; in the SPT experiment *f* (r,*t*) and *c_i_*(r,*t*) are the probabilities of finding the particle in the unbound and bound states, respectively.

In the Laplace space (*t* → *s*), the system of equations 1,2 can be reduced to a single equation with respect to the total concentration 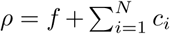 (In this paper, the original functions and their transforms can be distinguished by their arguments.):

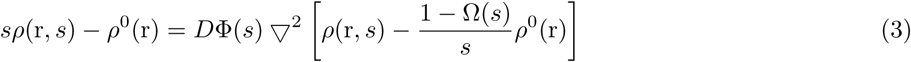

where

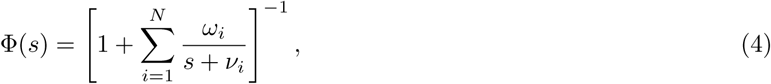

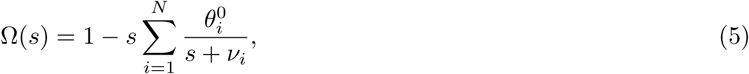

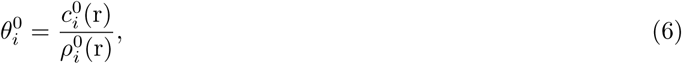

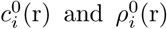 are the initial conditions. The solution of this equation corresponding to the initial condition *ρ*(r, *s*) = *δ_d_*(r) (propagator) has the form

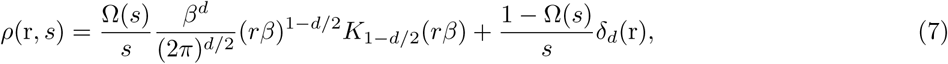

where *r* = |r|, *K*_1–*d*/2_(*r*) is the modified Bessel function of the second kind, *δ_d_*(r) is the *d*— dimensional delta-function, and 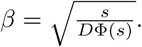

The multiple-trapping model is a generalization of the classical diffusion equation to the case when diffusing molecules can be delayed for a time by stationary binding sites. However, in many real disordered media, particularly in living cells, diffusion is also hampered by certain obstacles acting as potential barriers or as geometric constraints (1,5,6). The theoretical model, which claims to adequately describe the diffusion process in a medium with traps and obstacles, must take into account both these factors. In this paper, we propose a generalization of the multiple-trapping model that takes into account the presence of obstacles and provides some arguments indicating that the proposed model extends the descriptive capabilities of the multiple-trapping model.

## 2 Generalization of the multiple trapping model

In this section, we first show that the process described by the multiple-trapping model can be viewed as a subordinated stochastic process with the usual diffusion as the parent process and the renewal process as the leading process. Replacing in this subordinated process the usual diffusion into a process described by random barriers model, we obtain a generalization of the multiple-trapping model. Next, we show that the same result can be obtained within the framework of the lattice model. Some further generalizations and modifications are also considered.

Subordination is a mathematical method that broadens the applicability of classical transport models (7-11). In this method, the clock time of a stochastic process *X*(*t*) is randomized by introducing a new time *σ* = *S*(*t*). The resulting process *Y*(*t*) = *X*[*S*(*t*)] is said to be subordinated to the parent process *X*(*σ*), and *σ* is commonly referred to as the leading process or the operational time. If the processes *X*(*σ*) and *S*(*t*) are independent, then the propagator (one-point distribution function) of the resulting process is (12,11)

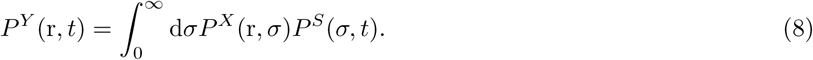

Here *P^X^*(r, *σ*) - one-point distribution function of the process *X*(*σ*), and *P^S^*(*t*, *σ*) - one-point distribution function of the process *S*(*t*). If the renewal process is taken as the leading process, then the propagator of the resulting process in the Laplace space (*t* → *s*) will be equal to (11)

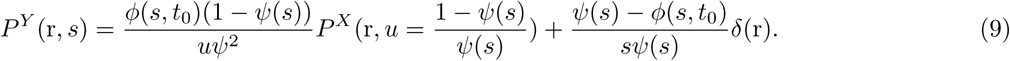

Here *P^X^*(r, *u*) is the Laplace image (*σ* → *u*) of the propagator of the parent process, *ψ*(*s*) is the Laplace image of the waiting time distribution function, *ϕ*(*s*, *t*_0_) is the recurrence time, *t*_0_ is the delay time. In the Laplace space (*t*_0_ → λ) the function *ϕ*(*s*, *t*_0_) is written as

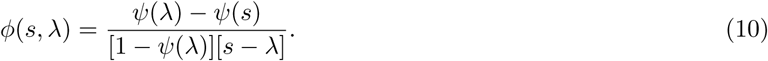

The delay time is the time elapsed between the start of the renewal process and the beginning of the process monitoring.

In order to obtain the propagator Eq. 7 by the subordination method, one must take the usual diffusion process (the Wiener process) as a parent process. The propagator of this process in the Laplace space (*σ* → *u*) is given by the formula

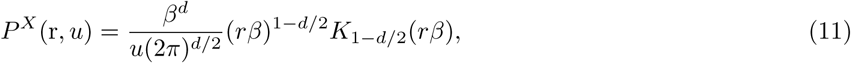

where 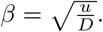 The distribution function *ψ*(*s*) should be given in the form

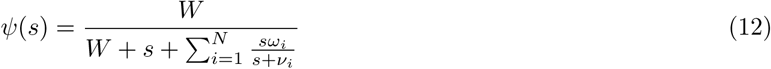

with the parameter *W* equal to one, and the recurrence time in the form *ϕ*(*s*, λ) = Ω(*s*, λ)*ψ*(*s*). The last relation will be consistent with relations Eq. 10 and Eq. 12, if the initial values 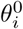 which appear in Eq. 5, are given in the form

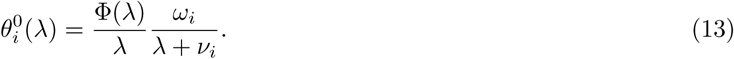

From this formula it follows that, with the delay time equal to zero, all 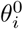 are equal to zero 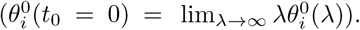 This means that the renewal process starts when the particle is in the transport state. At a delay time equal to infinity, all 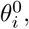 as it should, are equal to the equilibrium values 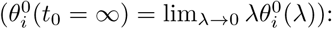

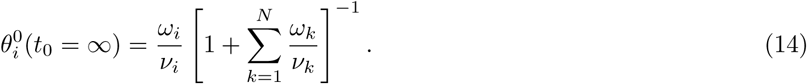

(Integrating Eqs. 1,2 over the whole space and solving them in the Laplace space *t* → λ under the initial conditions 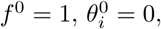 we find that the dependence of the probabilities *θ_i_* on the delay time is indeed described by Eq. 13 in the multiple-trapping model.)

Thus, the multiple-trapping model describes a subordinated stochastic process in which the role of the parent process is played by ordinary diffusion, and the role of the leading process is played by the renewal process with the waiting time distribution function Eq. 12. Such an interpretation of this model allows us to generalize it for the case when there are also barriers besides traps in a disordered medium. To do this, we take not ordinary diffusion as the parent process, but random walks in the random barriers model. The propagator of this model averaged over the ensemble of configurations satisfies the non-Markovian equation

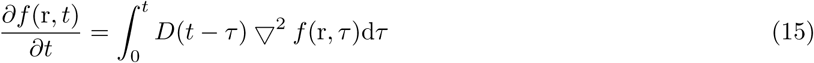

and in the Laplace space differs from the usual diffusion propagator Eq. 11 only in that the diffusion coefficient is a function of the Laplace variable: *D* = *D*(*u*). As a consequence, the propagator of the resulting process will have exactly the same form as the propagator (7):

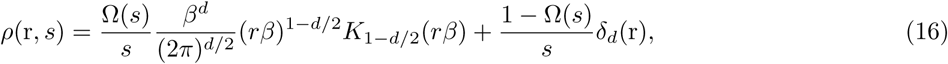

but with parameter *β* equal to 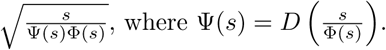 This propagator satisfies the equation

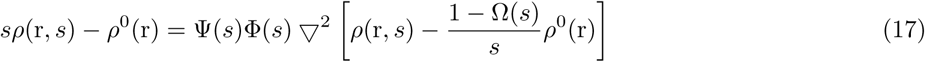

with the initial condition *ρ*^0^(r) = *δ_d_*(r). The analog of the system of equations 1,2 in this case is the system

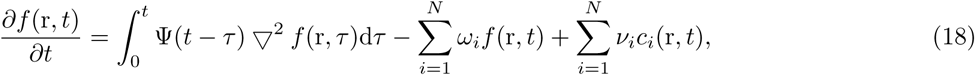

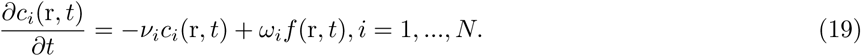

The equations of this system have the same meaning as equations 1,2. The difference between them is only in the transport term, which in this case is non-Markovian. This is due to the presence in this model of random barriers. It should be noted that it is impossible to obtain the system of equations 18,19 from the equation for the propagator of the model of random barriers Eq. 15 by simply adding in this equation the source terms and writing out the equations for the traps. This could be done if the Eq. 15 was Markovian. In the non-Markovian case, the memory function *D*(*t*) does not remain unchanged, but is modified, i.e. replaced by a function Ψ(*t*).

In the subordination method, it is assumed that the parent and leading processes are independent. In application to the model under consideration, this means that the traps and barriers are completely chaotically distributed. In the neighborhood to a barrier of any height, there can be a trap of any depth with equal probability. This is a limitation of this approach, since in real environments the distributions of traps and barriers can be correlated. To obtain more general models, we can use the Markov representation of the random barriers model at the mesoscopic level instead of the non-Markovian equation for the propagator Eq. 15 (13,14). In this approach, it is assumed that at each position in space the particle can be in M different transport states and that the probabilities of being in these states satisfy equations

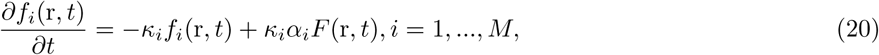

where *f_i_* is the probability of a particle being in the state of *i*-th type, *κ_i_* is the escape rate from the state of *i*-th type, *α_i_* fraction of states of *i*-th type, 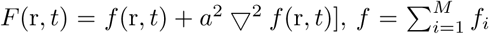 is the total probability, *a*^2^ is constant. Different states can be treated as localization’s of a particle surrounded by barriers of different heights. It follows from these equations that with an equiprobable initial state distribution 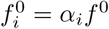 the total probability satisfies Eq. 15 with the memory function, which in the Laplace space has the form

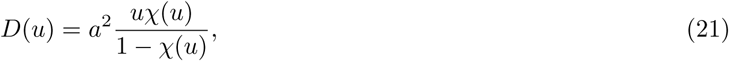

where 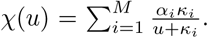 It should be noted that the equiprobable distribution 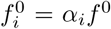 is also an equilibrium distribution for equations 20 (that is, equations 20 preserve the characteristic property of the random barriers model). Hence it follows that Eq. 15 is also valid for an equilibrium initial distribution. Since equations 20 are Markovian, one can add source terms to them without any additional modifications. Correlations can be taken into account here, i.e. in different equations one can add different source terms and write out different equations for the traps corresponding to these equations. To obtain the model considered above (Eqs. 18,19 or Eq. 17), it is necessary to add the same source terms to all equations 20 and to write out to each this equation the same set of equations for the traps:

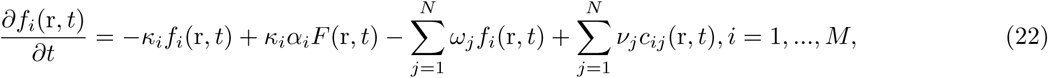

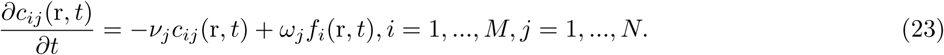

Here, the concentrations of particles in the traps have two indexes. One of them (*j*) corresponds to the trap type, and the other (*i*) – corresponds to the type of the transport state associated with this trap. It is assumed in this model that a particle can fall into a trap only from a transport state of one type and go from a trap to a transport state of the same type, i.e. a certain type of transport states corresponds to each trap. If we abandon this condition and assume that the transitions can occur into different states with the same probability, we get the model considered in (15):

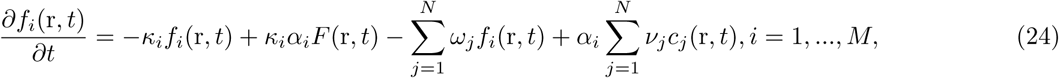

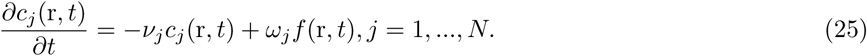

Both these systems of equations are brought to the same equation for the total concentration Eq. 17 (see Appendix A). In the first case, the memory function has the same form as in the method of subordination:

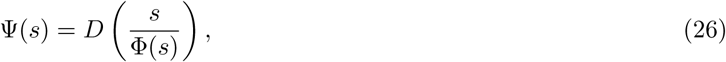

and in the second it is

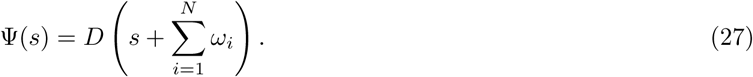

Here the memory function *D*(*u*) is given by Eq. 21. It’s easy to see that both these memory functions behave qualitatively as well as the function *D*(*u*), namely, they are positive monotonically increasing functions with negative second derivative (In the lattice model, which is the random barriers model, *D*(*u*) necessarily satisfies these properties (16,17)).

The simplest model with correlations has the form

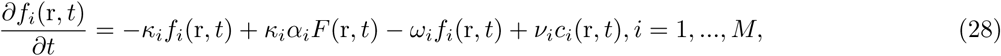

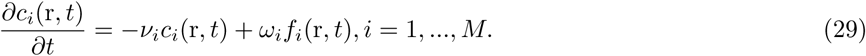

Here, each transport state is connected with its own type of traps, and with different states - different types of traps. In this model, the propagator has the same form as in the models without correlations (Eq. 17), but with other factors in front of the Bessel function and the delta function, as well as with another expression for the parameter *β* (see Appendix B).

In the following sections, the expression for the propagator Eq. 16 and equation 17 will be used to find theoretical expressions corresponding to the curves obtained in the experiments using fluorescence spectroscopy.

## 3 Fluorescence recovery after photobleaching

Together with the authors of paper (2), we consider the model that accounts for a finite nucleus and an arbitrary initial bleach profile. We assume a circular nucleus of radius *R_N_* that is photobleached at its center with an arbitrary, radially symmetric bleach profile *I*(*r*). Intensity measurements are made within a centered circle of radius *R_M_*.

The authors of (2) used the standard model of random traps (Eqs. 1,2) with one type of traps (*N* = 1). We will consider a more general model, taking into account the presence of barriers with an arbitrary number of types of traps (Eqs. 18,19 or Eq. 17). We’ll solve the problem in the Laplace space.

It is required to solve equation 17 in the two-dimensional region *r* ≤ *R_N_* with the initial condition *ρ*^0^(*r*) = *I*(*r*) and the boundary condition 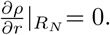 The solution should be finite at *r* = 0. Since the initial state of the system (before the flash) is in equilibrium, the values of 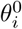 must be expressed by the formula Eq. 14. In this case, the function Ω(*s*) equals to 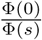 (This follows from Eq. 13 and Eq. 14). To obtain a curve corresponding to the experiment, it is necessary the obtained solution *ρ*(r, *s*) to integrate over the circle *r* ≤ *R_M_*.

The solution of equation 17 is represented in the form

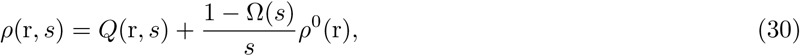

where the function *Q*(r, *s*) is a solution of equation

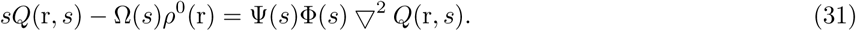

The solution of this equation can be expressed in terms of the Green’s function. In the case under consideration, the Green’s function can be defined as a bounded solution of the one-dimensional equation

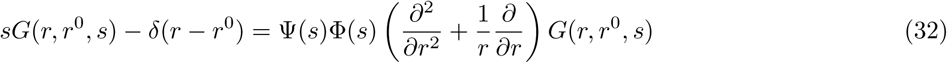

satisfying the boundary condition 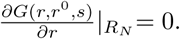 The function *Q*(r, *s*) is expressed as follows:

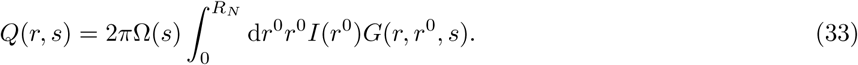

Integrating *ρ*(r, *s*) Eq. 30 over the circle *r* ≤ *R_M_*, we obtain relation

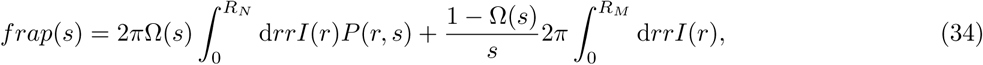

where

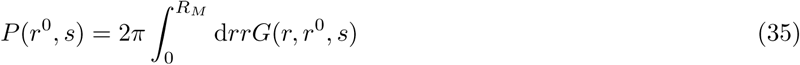

is a probability that the particle starting from the point *r*^0^ is in the circle *r* ≤ *R_M_*. This probability satisfies the equation analogous to Eq. 32, but with a different initial condition (18):

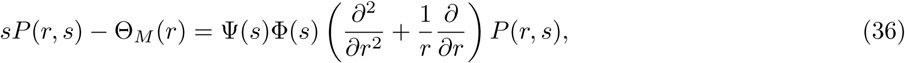

where Θ_*M*_(*r*) is the characteristic function of the interval 0 ≤ *r* ≤ *R_M_*. The boundary condition remains the same: 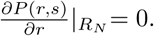

The bounded solution of equation 36 has the form

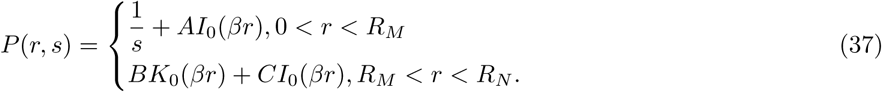

Here *I*_0_ and *K*_0_ are the modified Bessel functions of the first and second kind, respectively, 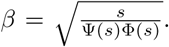 The coefficients *A*, *B* and *C* are found from the boundary condition and the requirement of continuity of the function *P*(*r*, *s*) and its first derivative at the point *r* = *R_M_*. The final expression for the function *P*(*r*, *s*) is written as

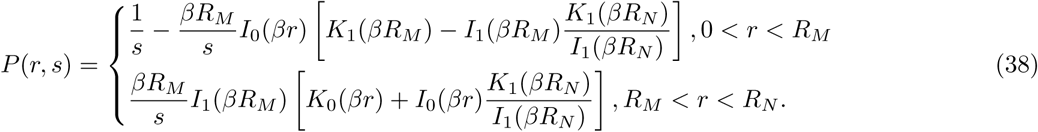

(In the course of the calculations, the following relations between the modified Bessel functions were used: 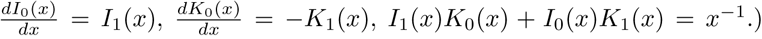 Substituting this expression in Eq. 34, we obtain the final expression for *frap*(*s*):

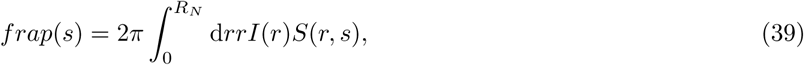

where the function *S*(*r*, *s*) has the form

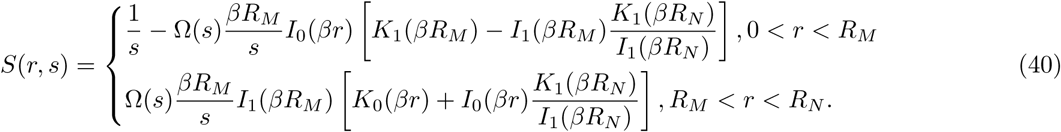

In the case of infinite nucleus and special profile *I*(*r*) = 0 for 0 < *r* < *R_M_* and 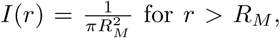 we

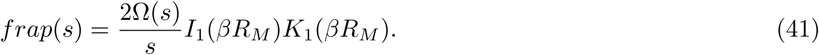

(In the course of the calculations, the following properties of the modified Bessel functions were used: the quotient 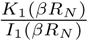 is equal to zero for infinite *R_N_*; the integral 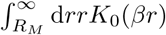 is equal to 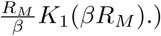 This formula was obtained in papers (3,4) for the case *N* = 1. In these papers, Φ(*s*) was equal to 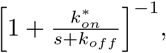 and the function Ω(*s*), respectively, was 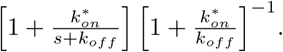. The function Ψ(*s*) in the paper (3) was equal to 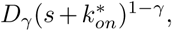 and in the paper (4) it was equal to the constant *D_f_*.

The transition to real time in Eq. 39 must be carried out numerically. In some cases, one can get an analytical expression in the form of an infinite series. To do this, we expand the function Eq. 40 in a Fourier series with respect to the Bessel functions *J*_0_(*α_i_r*) on the interval (0, *R_N_*) and perform term-by-term integration in Eq. 39. The result is an expression

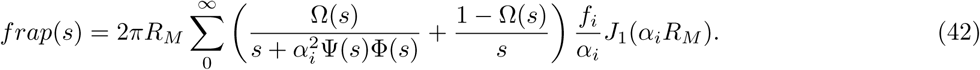

Here 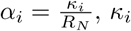 are the zeros of the function *J*_1_(*r*),

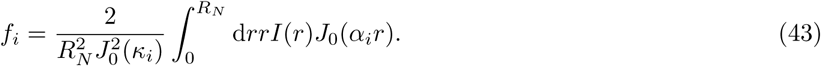

(This result can also be obtained by solving the initial problem by the method of separation of variables.) For functions Φ(*s*) and Ψ(*s*) of simple form, each term of this series can be inverted analytically. In particular, when 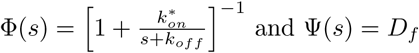 we obtain the result obtained earlier in paper (2).

## 4 Fluorescence correlation spectroscopy

Usually, the FCS curve is represented as an integral from the product of the propagator to the apparatus function that determines the laser intensity distribution (19):

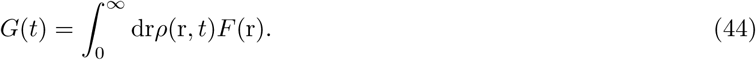

If the laser intensity distribution is Gaussian, then the apparatus function is written as

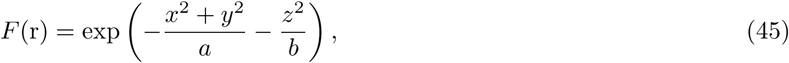

where *a* and *b* are positive parameters. In the case of normal diffusion, the integral Eq. 44 is calculated explicitly:

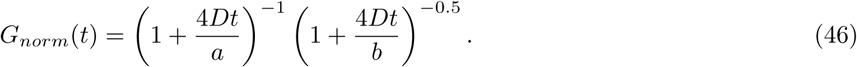

Using this result, we find the Laplace image of the FCS curve for the model considered here. In the Laplace space, Eq. 44 can be written in the form

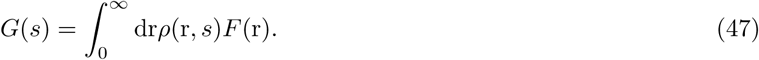

In the model under consideration, the propagator *ρ*(r, *s*) is given by Eq. 16. The dependence on r of the first term on the right-hand side of this formula does not differ from the dependence on r of the diffusion propagator Eq. 11. Consequently, the integral of this term will have the same functional form as the integral of the diffusion propagator. The difference will be only in the appearance of a factor Ω(*s*) and in replacing the diffusion constant *D* by the product Ψ(*s*)Φ(*s*). The second term is integrated trivially. Taking into account that for normal diffusion the Laplace image of the FCS curve has the form

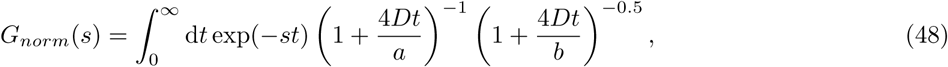

we find the Laplace image of the FCS curve for the model under consideration:

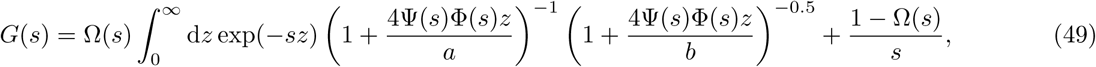

Since the FCS experiment is performed under equilibrium conditions, the function Φ(*s*) in this formula is equal to 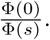 To avoid misunderstandings, we renamed the integration variable *t* (*t* → *z*) because in this case it is not a time.

## 5 Single particle tracking

SPT experiment consists in tracing the trajectory of an individual particle. A labeled particle is introduced into the test sample and monitored with a video microscope. The trajectory of the particle *r*(*t*) is recorded for a time sufficient to have a complete statistics of spatial displacements. The information thus obtained allows, in principle, to find all the correlation functions. In particular, the propagator Eq.16 and all the quantities calculated with it can be found. We give the expression for the mean square displacement:

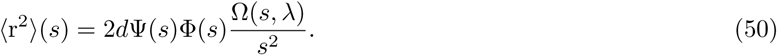

Here, the dependence of the initial probabilities Eq. 13 on λ is taken into account, which makes it possible to calculate such a widely used quantity as the mean-square displacement averaged over the ensemble and in time (delay time). In the case of a fully equilibrated system (when 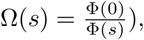 Eq. 50 reduces to

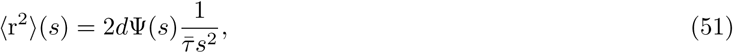

where 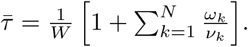 The characteristic feature of this expression is that it does not depend on the function λ(*s*); therefore, if there are no barriers, i.e. if Ψ(*s*) does not depend on *s*, this mean-square displacement is a linear function of time.

To go to the real time in Eq. 50, we rewrite it in the form

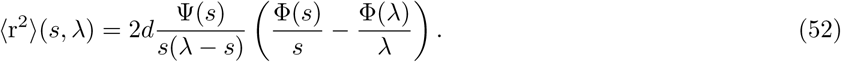

Performing inverse Laplace transforms with respect to *s* and λ, we obtain the expression for the mean square displacement of particles as a function of time *t* and delay time *t*_0_:

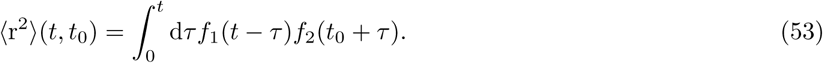

where *f*_1_(*t*) is the original of the function 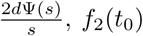 is the original of the function 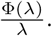 After averaging this expression with respect to *t*_0_, we obtain the expression for the mean square displacement of the particles, averaged over both time and ensemble (here the notation for the variable *t* (*t* → Δ) is changed):

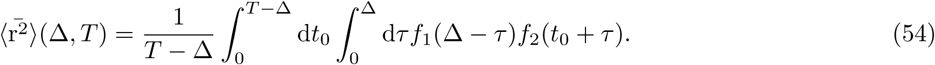

Here, *T* – Δ is the period of time over which the averaging is carried out. If the condition Δ << *T* is fulfilled, then by means of the substitutions *T* – Δ → *T* and *t*_0_ + *τ* → *t*_0_ the last expression can be reduced to the form

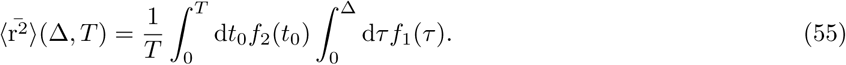

## 6 Discussion and conclusion

The model considered in this paper has two functions as parameters: Φ(*s*) and Ψ(*s*). The form of the function Φ(*s*) is given by equation 4. For its unambiguous determination, it is necessary to specify the parameter *N* and 2*N* of the parameters *ν_i_* and *ω_i_*. The function Ψ(*s*) is expressed in terms of the indeterminate function *D*(*s*), therefore its form can be set to a considerable extent arbitrarily. It is only necessary that it be a positive monotonically increasing function with negative second derivative. As the simplest functions satisfying these conditions, one can take a power function Ψ(*s*) = *A*(*s* + *k*)^*n*^, where *A* > 0, *k* > 0, 1 > *n* > 0 or a fractional rational function 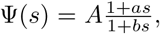 where *A* > 0, *b* > 0, *a* > *b*. If it is assumed that there is only one type of binding centers, *N* = 1, then the model will contain five parameters: *ν_i_*, *ω_i_*, *A*, *k*, *n* in the first case and *ν_i_*, *ω_i_*, *A*, *a*, *b* in the second case. The numerical values of these parameters can be found by the method of least squares. The input data can be the FRAP curve, the FCS curve, the propagator, the mean square displacement. The corresponding theoretical expressions are: Eqs.39,40 for the FRAP curve, Eq. 49 for the FCS curve, Eq. 16 for the propagator, Eq. 54 for the mean square displacement. If there is an experimental dependence of the mean square displacement on time, the function Ψ(*s*) can be found up to a multiplicative constant, independently of Φ(*s*), using equation 51. Having found the functions Φ(*s*) and Ψ(*s*), we know the parameters *ν_i_* and *ω_i_*, which characterize the binding properties of biomolecules. In order to establish the diffusion properties, one of the functional equations 26 or 27 with respect to the function *D*(*s*) must be additionally solved. Which of these equations must be solved depends on which of the two systems of equations 22,23 or 24,25 is more adequate in the case under consideration. However, in order to clarify this, it is necessary to have some information about the structure of the disordered medium under consideration.

Here are two examples that confirm the correctness of the proposed model. The first example. The authors of paper (3) used a special case of the model in question to describe the experimental data obtained in paper (4). It was shown that a model with a function Ψ(*s*) different from a constant is more adequate than the traditional model. It describes the experimental data for different sizes of the photobleached volume with the same set of parameters. It should be noted that the authors of (3) interpreted their equations differently than it is done in this paper. They believed that the appearance of the function Ψ(*s*) is due to the anomalous diffusion described by the continuous-time random walk model (CTRW). By the fact that in the framework of this model equation 15 is valid only in the nonequilibrium case, they have neglected.

The second example. The model under consideration gives the same result as the CTRW on fractals (20). Namely, if the function *D*(*s*) has the form

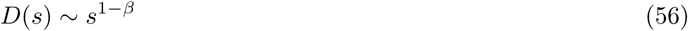

(i.e. it describes the anomalous subdiffusion due to barriers), and the function Φ(*s*) has the form

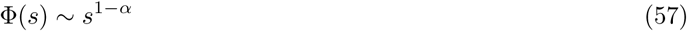

(i.e. it describes the anomalous subdiffusion due to the traps), then the function Ψ(*s*) Eq. 26 will look like

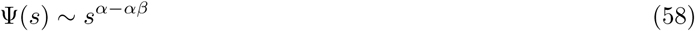

and the time-averaged mean square displacement Eq. 55 is written in the form

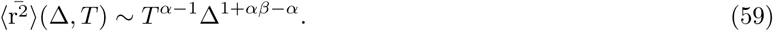

The same dependence was obtained experimentally in paper (5).

In conclusion, in this paper we propose a mathematical model for describing experiments using fluorescent microscopy in disordered media containing binding sites and obstacles. In the Laplace space, the expressions for the propagator, for the mean square displacement as a function of the observation time and delay time, for the FRAP curve and for the FCS curve are derived. The model can be used to find parameters characterizing the diffusion and binding properties of biomolecules in living cells.

## 7 Appendix A

In the Laplace space, the equations 22,23 have the form

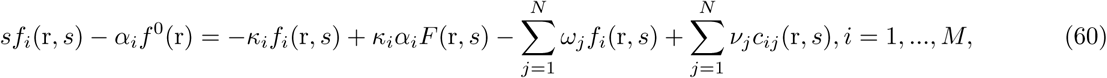

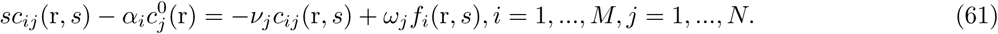

We express from the second equation *c_ij_*:

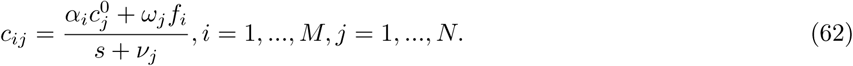

We substitute this expression in the first equation:

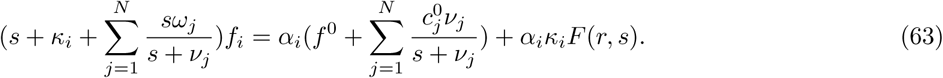

We express from here *f_i_* and sum over *i*:

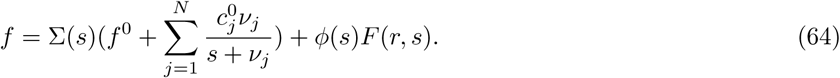

Here

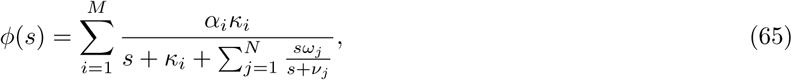

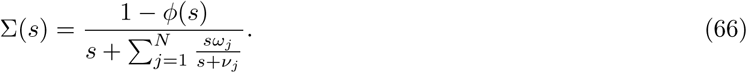

By dividing Eq. 64 by Σ(*s*) and transforming, we obtain

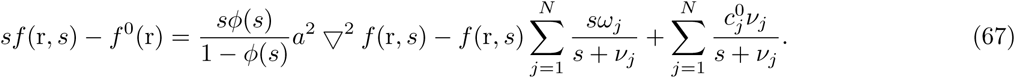

Further, summing equations 61 with respect to *i* and *j*, and adding to Eq. 67, we obtain

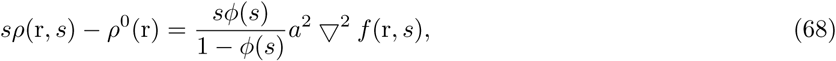

where 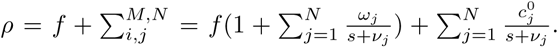 Expressing *f* in this equation in terms of *ρ*, we finally obtain equation 17 with function Ψ(*s*) equal to Eq. 26.

In the Laplace space, equations 24,25 have the form

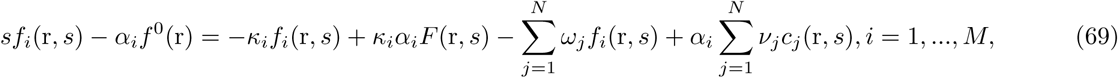

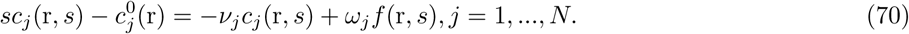

From these equations, as a result of calculations similar to the previous ones, we obtain equation 17 with the function tfΨ(*s*) equal to Eq. 27.

## 8 Appendix B

In the Fourier-Laplace space, equations 28,29 have the form

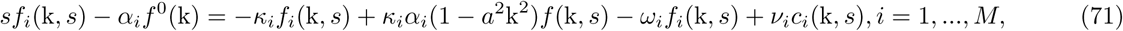

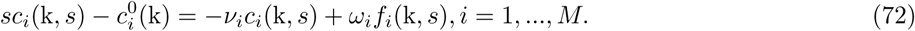

We assume that at the initial instant of time the whole probability is concentrated at one point *r* = 0 and, therefore, the initial values 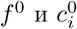 do not depend on *k*. We express from the second equation *c_i_*:

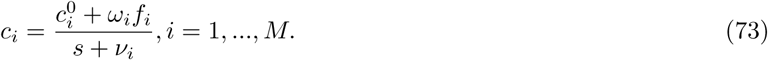

We substitute this expression in the first equation and solve it with respect to *f_i_*:

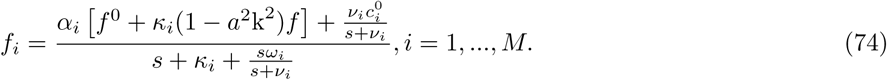

We summarize this relation with respect to *i* and express *f*:

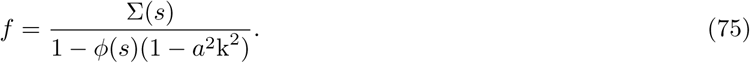

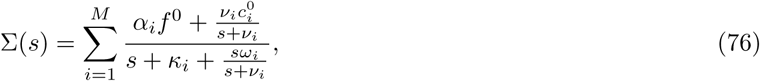

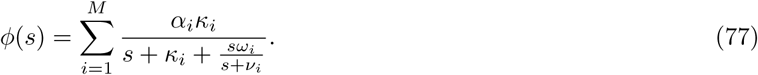

The expression (1 – *a*^2^*k*^2^)*f* can be transformed to the form 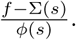 Using expressions obtained, we find propagator 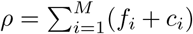 in the Fourier-Laplace space. Performing the inverse Fourier transform, we obtain an expression similar to Eq. 17 but with other factors in front of the Bessel function and the delta function, as well as with another expression for the parameter *β*.

We note an interesting property of this model. If the initial distribution over traps of different types is equilibrium, 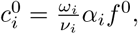 then the function Σ(*s*) takes the form

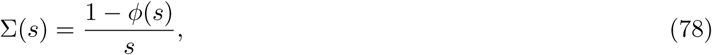

and the expression for the probability of being in the mobile state Eq. 75 turns into

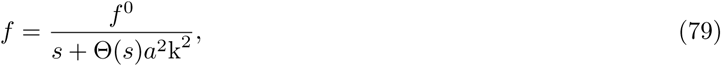

where 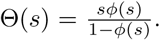 This expression coincides with the expression for the propagator in the continual version of the CTRW model.

